# Sequencing of *Panax notoginseng* genome reveals genes involved in disease resistance and ginsenoside biosynthesis

**DOI:** 10.1101/362046

**Authors:** Guangyi Fan, Yuanyuan Fu, Binrui Yang, Minghua Liu, He Zhang, Xinming Liang, Chengcheng Shi, Kailong Ma, Jiahao Wang, Weiqing Liu, Libin Shao, Chen Huang, Min Guo, Jing Cai, Andrew KC Wong, Cheuk-Wing Li, Dennis Zhuang, Ke-Ji Chen, Wei-Hong Cong, Xiao Sun, Wenbin Chen, Xun Xu, Stephen Kwok-Wing Tsui, Xin Liu, Simon Ming-Yuen Lee

## Abstract

*Panax notoginseng* is a traditional Chinese herb with high medicinal and economic value. There has been considerable research on the pharmacological activities of ginsenosides contained in *Panax* spp.; however, very little is known about the ginsenoside biosynthetic pathway. We reported the first *de novo* genome of 2.36 Gb of sequences from *P. notoginseng* with 35,451 protein-encoding genes. Compared to other plants, we found notable gene family contraction of disease-resistance genes in *P. notoginseng*, but notable expansion for several ATP-binding cassette (ABC) transporter subfamilies, such as the Gpdr subfamily, indicating that ABCs might be an additional mechanism for the plant to cope with biotic stress. Combining eight transcriptomes of roots and aerial parts, we identified several key genes, their transcription factor binding sites and all their family members involved in the synthesis pathway of ginsenosides in *P. notoginseng*, including dammarenediol synthase, *CYP716* and *UGT71*. The complete genome analysis of *P. notoginseng*, the first in genus *Panax*, will serve as an important reference sequence for improving breeding and cultivation of this important nutraceutical and medicinal but vulnerable plant species.

## Introduction

Genus *Panax* in the Araliaceae, contains some medicinally and economically important ginseng species including *P. ginseng*, *P. quinquefolius* and *P. notoginseng* (Sanqi in Chinese)^1^. Because *P. notoginseng* is an important Chinese traditional medicine, two draft genomes of *P. notoginseng* were published in 2017. Chen etc.’s genome assembly was about 2.39 Gb with a contig N50 of 16 kb and scaffold N50 of 96 kb and Zhang etc.’s was a ~1.85 Gb assembly with scaffold N50 of 158 kb and contig N50 of 13.2 kb^2,3^. There was a notable difference of genome assembly size (~540 Mb) between two genomes. In general, the estimation of genome size using flow cytometry analysis is considered as a golden standard approach and the estimated result of diploid P. *notoginseng* was about 2.31 Gb^3^. Besides, these two papers also annotated the repetitive sequences and protein-coding genes. Chen etc. annotated ~75.94% repetitive sequences (~1.71 Gb) and 36,790 genes, and Zhang etc. annotated 61.31% repetitive sequences (~1.13 Gb) and 34,369 genes in the assembly, respectively. Zhang etc.’s genome has a longer scaffold N50 size, but shorter contig N50 size, total length and proportion of repetitive sequences comparing to Chen etc.’s genome, suggesting Zhang etc.’s assembly might miss a large amount of repetitive sequences. Thus, due to the high proportion of repetitive sequences in this species, it is a big challenge to obtain a good assembly for this complex genome.

*Panax notoginseng* is susceptible to a wide range of pathogens and identification of the genes conferring disease resistance has been a major focus of research^4^, which greatly reduces its economic benefits. The main class of disease-resistance genes (R-genes) consist of a NBS, a C-terminal LRR and a putative coiled coil domain (CC) at the N-terminus or a N-terminal domain with homology to the mammalian toll interleukin 1 receptor (TIR) domain^5^. Generally, the LRR domain is often involved in protein–protein interactions, with an important role in recognition specificity as well as ligand binding^6^. Besides, ATP-binding cassette (ABC) proteins have also been reported to play a crucial role in the regulation of resistance processes in plants^7^, most of which modulate the activity of heterologous channels or have intrinsic channel activity. The reason underlying the poor resistance of *P. notoginseng* is still unclear. Saponin constituents, also known as ginsenosides, are the major active ingredients in genus *Panax*^8,9^ and have diverse pharmacological activities, such as hepatoprotection, renoprotection, estrogen-like activities and protection against cerebro-cardiovascular ischemia and dyslipidaemia in experimental models^10-14^. Xuesaitong (in Chinese) is a prescription botanical drug, manufactured from saponins in the root of *P. notoginseng*^10^, used for the prevention and treatment of cardiovascular diseases, and is among the top best-selling prescribed Chinese medicines in China^15,16^. Compound Danshen Dripping Pill, a prescription botanical drug for cardiovascular disease^17^, comprises *P. notoginseng*, *Salvia miltiorrhiza* and synthetic borneol, and has successfully completed Phase II and is undergoing Phase III clinical trials in the United States. Although different ginseng species have been one of the most important traded herbs for human health around the world, the industry is facing elevating challenges, such as increasing disease vulnerability of the cultivated plant and an as yet undetermined disease preventing continuous cultivation on the same land, leading to a trend of decreasing production^18^.

Plant metabolites play very important roles in plant defense mechanisms against stress and disease, and some have important nutritional value for human consumption. For instance, capsaicinoids and caffeine present in pepper and coffee, respectively, have important health benefits. The recently completed whole-genome sequencing of coffee and pepper provide new clues in understanding evolution and transcriptional control of the biosynthesis pathway of these metabolites^19,20^. Ginsenosides, the oligosaccharide glycosides of a series of dammarane- or oleanane-type triterpenoid glycosides^9^, are a group of secondary metabolites of isoprenoidal natural products, specifically present in genus *Panax*. Based on the structure differentiation of the sapogenins (aglycones of saponin), dammarane-type ginsenosides can be classified into two subtypes: protopanaxadiol (PPD) and protopanaxatriol (PPT) types^9^. Up to now, over 100 structurally diversified ginsenosides have been isolated. Interestingly, the two types (PPD and PPT) of ginsenosides have been reported to exhibit opposing biological activities; for instance, pro-angiogenesis and anti-angiogenesis^21,22^. The whole plant of *P. notoginseng* contains both PPD- and PPT-type ginsenosides but the aerial parts (e.g. leaf and flower) contain a higher abundance of PPD compared to roots^23,24^. Herein, we characterize the genome of *P. notoginseng*, the population of which has a relatively uniform genetic background. This was aimed to determine the genetic causes of weakened resistance of *P. notoginseng* as well as to understand evolution and regulation of the triterpenoid saponin biosynthesis pathway leading to the production of specialized ginsenosides in genus *Panax*.

## Results

### Genome assembly and annotation

We *de novo* sequenced and assembled the *P. notoginseng* genome from Wenshan City of China using ~256.07 Gb data (~104.09-fold) generated by Illumina HiSeq2000 and ~13 Gb (~6-fold, average read length of ~9kb) SMRT sequence data generated by Pacbio RSII sequencing system (**Supplementary Table 1**). The final genome assembly spanned about 2.36 Gb (~95.93% of the estimated genome size) with scaffold N50 of 72.37 kb and contig N50 of 16.42 kb (**Supplementary Table 2**, **3** and **Supplementary Fig. 1**). To evaluate our assembly, we firstly checked the links within scaffolds based on the pair-end relationships of sequencing reads and the result revealed that pair-end information supported the links of scaffolds well (**Supplementary Fig. 2**). Secondly, we obtained the unmapped sequencing reads against our genome assembly and re-assembled them into ~359 Mb sequences which contained ~95.05% repetitive sequences (**Supplementary Table 4**). Using KAT sect tool^25^, we also calculated the sequencing depth distribution of flanking sequences of gap regions, we observed that ~71% of sequences nearby junction regions have higher sequencing depth (**Supplementary Fig. 3**), indicating that majority of the gap regions were repetitive sequences. Besides, leaves and roots of *P. notoginseng* were collected from 1-, 2- and 3-year-old *P. notoginseng* and flowers collected from 2- and 3-year-old *P. notoginseng*. An average of ~6.06 Gb raw data were obtained from each sample sequenced using Illumina HiSeq 2000 (**Supplementary Table 5**). More than 95.42% of transcripts (coverage ratio >= 90%) could be unambiguously mapped into assembly sequences revealing the good integrity of genome assembly (**Supplementary Table 6**). Of the genome, 72.45% of repetitive sequences ratio was estimated by Jellyfish^26^ but only 1.24 Gb (51.03%) was found to be repetitive sequences, which is more than that of carrot genome (~46%)^27^, with long terminal repeats the most abundant (~95.08% of transposable elements) (**Supplementary Table 7**). We totally predicted 35,451 protein-encoding genes in the *P. notoginseng* genome by integrating the evidences of *ab initio* prediction, homologous alignment and transcripts (**Supplementary Table 8**). Assessing the *P. notoginseng* genome assembly and annotation completeness with single-copy orthologs approach (BUSCO)^28^ showed that 895 (93%) complete single-copy genes containing 246 (25%) complete duplicated genes were validated, similar to that of cotton^29^ and *Dendrobium officinale*^30^. In combination with the RNA-seq mapping results above, it was unambiguous that our gene set was available for further analysis.

### Genome evolution of *Panax* species

*Panax notoginseng* is the first sequenced genome of genus *Panax*, thus we compared its genome to other related species to reveal genome evolution of *Panax* species. To analyze *P. notoginseng* evolution, we first constructed the phylogenetic tree using published closely-related species (**Supplementary Fig. 4**). We estimated the species divergence time between *P. notoginseng* and *D. carota* to be about 71.9 million years ago (**Supplementary Fig. 4**). Compared to closely related species (*D. carota*, *S. tuberosum*, *S. lycopersicum* and *C. annuum*), we found 1,144 unique gene families (containing 2,714 genes) with no orthologs in other species (**Supplementary Fig. 5**), which were possibly related to specific features of *P. notoginseng*. Thus, we performed functional enrichment of these genes to find several interesting Gene Ontology (GO) terms likely related to saponin biosynthesis, such as UDP-glucosyltransferase (UGT) activity (P = 0.003, **Supplementary Fig. 6**). To determine features and specific functions in *P. notoginseng*, we identified a total of 1,520 expanded gene families (**Supplementary Fig. 4**) and further performed Kyoto Encyclopedia of Genes and Genomes (KEGG) and GO functional enrichment analyses for these genes. We found several significant gene families related to saponin biosynthesis, such as sesquiterpenoid and triterpenoid biosynthesis (P = 0.006, **Supplementary Table 9**).

### Resistance genes

Most cloned R-genes in the plant encode nucleotide-binding site and leucine-rich-repeat (NBS-LRR) domains. In our *P. notoginseng* genome assembly, 129 NBS-LRR-encoding genes were detected, which was notably fewer than *Arabidopsis thaliana* (183), *D. carota* (170), *S. tuberosum* (441), *S. lycopersicum* (282) and *C. annuum* (778), and similar to that of *C. sativus* (89)^31^ (**Supplementary Table 10**). Considering R genes were known to be small and in repetitive areas and our assembly with much gaps in the repetitive regions, the total number of R gene of *P. notoginseng* would be likely to underestimated. However, the number of genes in the TIR_NBA_LRR subfamily was still relatively expanded according to the phylogenetic analysis using *P. notoginseng*, tomato and carrot, which possibly be related to the resistance character of *P. notoginseng* (**Fig. 1a** and **1b**).

**Fig. 1.**
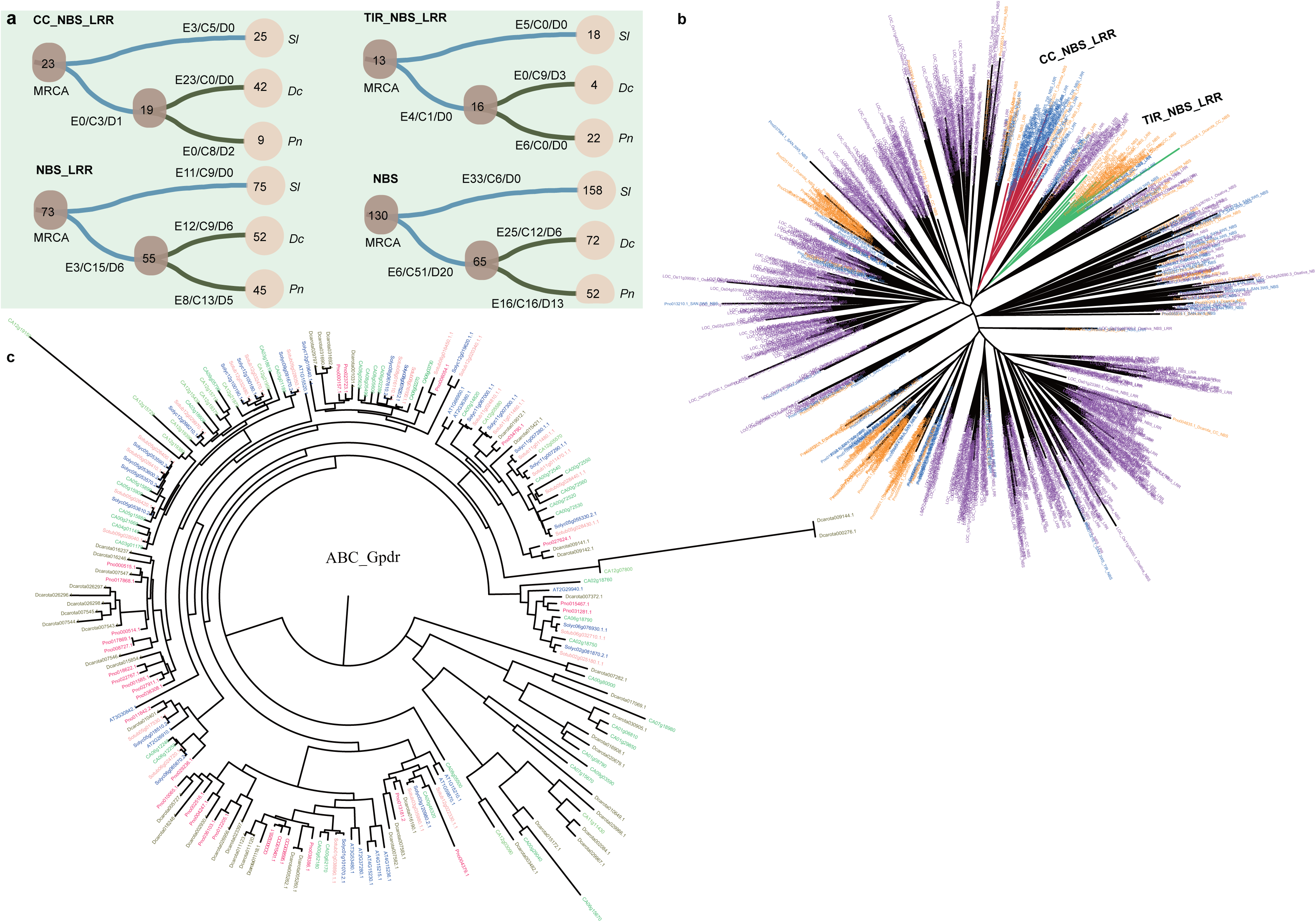
The gene families related to resistance of *P. notoginseng*. **(a)** The evolution of R-genes with NBS domains among *S. lycopersicum* (*Sl*), *D. carota* (*Dc*) and *P. notoginseng* (*Pn*). The numbers in circles represent the gene numbers of a species and the numbers in clades represent the gene number of expanded gene families (E), contracted gene families (C) and extinct gene families (D). NBS represents a gene that only contains the NBS domain, and NBS_LRR, TIR_NBS_LRR and CC_NBS_LRR follow a similar representation. **(b)** Phylogenetic tree of R genes of *P. notoginseng, D. carota* and *Oryza sativa* (outgroup). Blue label is *P. notoginseng*, orange is *D. carota* and purple is *Oryza sativa*. **(c)** The gene trees of ABC Gpdr subfamily of *P. notoginseng* and other five plants. The red represents genes of *P. notoginseng*, the green is pepper, the orange is potato, dark blue is tomato, the cyan is carrot and the blue is Arabidopsis.

### ABC transporter gene family

We identified a total of 153 ATP-binding cassette (*ABC*) genes in *P. notoginseng*, which were classified into nine subfamilies (**Supplementary Table 11** and **Supplementary Fig. 7**). Generally, compared with *A. thaliana* (12), *Daucus carota* (5), *Solanum lycopersicum* (10), *S. tuberosum* (13) and *Capsicum annuum* (10), there were notably fewer members of subfamily A in *P. notoginseng* (3); whereas, there were notably more of subfamily B in *P. notoginseng* (28, 28, 33, 38, 34 and 47 for subfamily B, respectively) (**Supplementary Table 11**). In detail, two genes were notably contracted for subfamily A (**Supplementary Fig. 8**) – a subfamily reportedly involved in cellular lipid transport^32^.

For subfamily B, we found a specific expanded clade in *P. notoginseng* containing 11 genes, three duplication expansions and one contraction (**Supplementary Fig. 7** and **Supplementary Fig. 9**). In particular, duplication expansions were Pno037415.1 and Pno028859.1 (AT2G36910.1 homologs); Pno009402.1 and Pno037963.1 (AT3G28860.1 homologs); and Pno034123.1 and Pno018232.1 (AT5G39040.1 homologs). One contraction was Pno003332.1 (AT4G28620.1, AT4G28630.1 and AT5G58270.1 homolog). Interestingly, the expanded genes have been reported as involved in aluminum resistance and auxin transport in stems and roots, respectively^33,34^. The contracted gene was found to be involved in iron export^35^. In the Gpdr subfamily, we also found a notable expansion (*P. notoginseng*: 10 vs. *A. thaliana*: 1, AT1G15520.1) (**Fig. 1c**). The homolog of AT1G15520.1 is known to be related to pleiotropic drug resistance and abscisic acid (ABA) uptake transport^36,37^.

### Key genes involved in ginsenoside biosynthesis of *P. notoginseng*

#### Dammarenediol synthase

Ginsenosides are committed to be synthesized from dammarenediol-II after hydroxylation via cytochrome P450 (*CYP450*)^38^ and then glycosylation by glycosyltransferase (*GT*)^39^. Dammarenediol-II is synthesized from farnesyl-PP, which is synthesized via the terpenoid backbone biosynthesis pathway. There are three enzymes involved from farnesyl-PP to dammarenediol-II: farnesyl-diphosphate farnesyltransferase (*FDFT1*, K00801), squalene monooxygenase (*SQLE*, K00511) and dammarenediol synthase (*DDS*, K15817) (**Fig. 2a**). In total, we identified three *SQLE*, two *FDFT1* and one *DDS* in *P. notoginseng*. Combining with the results of real-time PCR, *DDS* (Pno034035.1) was found with extensive higher expression in the root than aerial parts and in 3-year old plant both root and aerial parts were found with higher expression of *DDS* (Pno034035.1). We also found that expressions of one *FDFT1* (Pno029444.1) and one *SQLE* (Pno022472.1) were higher in roots compared with flowers and leaves, and one *SQLE* (Pno009162) gene showed the opposite pattern according to RNA-seq, while the real-time PCR results suggested an increasing expression year after year of all these three genes (**Fig. 2b**). Phylogenetic analysis showed that *DDS* of *P. notoginseng* and *P. ginseng* had a very high (98.96%) amino acid sequence identity, differing in only eight amino acids, and with fewer genes than other species (**Supplementary Fig. 10**). We further found there were three amino-acid residue insertions (L194, A195 and E196) in the *DDS* sequences of *P. notoginseng* and *P. ginseng*, which were located in the cyclase-N domain (**Fig. 2c**). We used the SWISS-MODEL^40^ to conduct structural modeling of *DDS* protein using the human oxidosqualene cyclase (*OSC*) in complex with Ro 48-80771^41^(SMTL: 1w6j.1) as the template. When the locations of the amino acids were mapped to the protein 3D structure with amino-acid identity of 40%, two residue insertions (L194 and A195) were located between two DNA helices, which are the presumed substrate-binding pocket and the catalytic residues (**Fig. 2d**). These three amino acids of *DDS* sequences of *P. notoginseng* probably played an important role in the Dammarenediol-II biosynthesis.

**Fig. 2.**
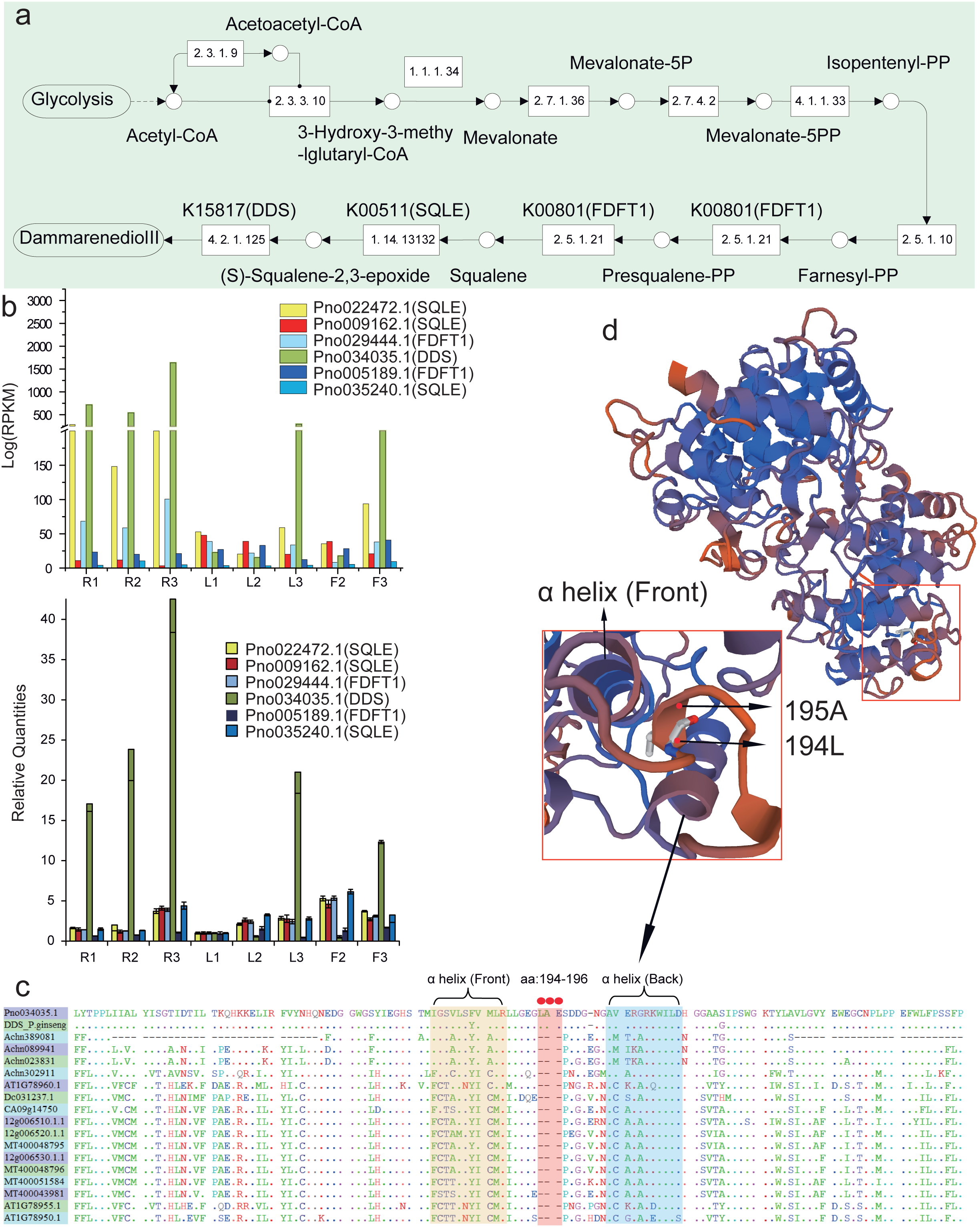
The preliminary steps of ginsenoside biosynthesis and several key gene families. **(a)** The dammarenediol-II synthesis pathway from glycolysis involves terpenoid backbone biosynthesis and finally catalysis by DDS. **(b)** Comparison of gene expressions of three key enzyme gene families between the roots and aerial parts (flowers and leaves) and the qPCR results. R1 represents one-year-old roots of *P. notoginseng*, R2: two-year-old roots, R3: three-year-old roots from plant B, L1: one-year-old leaves, L2, two-year-old leaves, L3: three-year-old leaves, F2: two-year-old flowers, F3: three-year-old flowers. **(c)** Protein sequence multiple comparisons of DDS among *P. notoginseng*, *P. ginseng* and other representative plants. Stars represent amino acids conserved in all protein sequences, red circles represent amino acids that are the same in *P. notoginseng* and *P. ginseng*, but missing in all other plants. **(d)** The protein 3D structure of DDS of *P. notoginseng* constructed by SWISS-MODEL. The arrows indicate two amino acids specific to *P. notoginseng* and *P. ginseng*.

#### CYP450s

*CYP450*s and *GT*s have been demonstrated to be involved in hydroxylation or glycosylation of aglycones for triterpene saponin biosynthesis. We identified a total of 268 *CYP450* genes in *P. notoginseng* using 243 *CYP450* homolog genes of *A. thaliana*, which is consistent with the report that there are around 300 *CYP450* genes in genomes of flowering plants^42^. To compare with other species, we also identified *CYP450* genes of four other genomes—*D. carota* (325), *S. tuberosum* (465), *S. lycopersicum* (252) and *C. annuum* (256)—using the same parameters as for *P. notoginseng* (**Supplementary Table 12** and **Supplementary Fig. 11**). We further classified the subfamilies of *CYP450* genes for each species based on the category of *CYP450* in *A. thaliana* and nine *CYP716* genes were identified in *P. notoginseng* (**Fig. 3a**). *CYP716A47* was reported to produce PPD^43^ and *CYP716A53v2* was reported to be involved in the synthesis of PPT in *P. ginseng* (**Fig. 3b**). Moreover, considering the role of different types of notoginsenoside in different parts/tissues of *P. notoginseng*— that is, PPT was relatively higher in roots while PPD was higher in aerial parts (leaves and flowers) (**Fig. 3c**)— we sequenced the transcriptomes of leaves, roots and flowers collected from different ages of *P. notoginseng*. We found Pno012347.1 (*CYP716A47* homolog), Pno002960.1 (*CYP716A53v2* homolog) and Pno011760.1 were always more highly expressed in roots than in leaves and flowers for all ages; while Pno021283.1 (*CYP716A47* homolog) was the opposite (**Fig. 3d**). Thus, Pno012347.1 and Pno021283.1 were the candidate genes involved in PPD biosynthesis and Pno002960.1 was the candidate gene involved in PPT biosynthesis in *P. notoginseng*. The expression levels of three *CYP716* genes (Pno012347.1, Pno002960.1 and Pno011760.1) validated using real-time PCR were consistent with RNA-seq. Meanwhile, we identified another 22 *CYP450* genes (including *CYP78*, *CYP71*, *CYP72* and *CYP86*), which had the same expression pattern as three *CYP716A53v2*-homolog genes (**Fig. 3d**), indicated they may be involved into PPT biosynthesis.

**Fig. 3.**
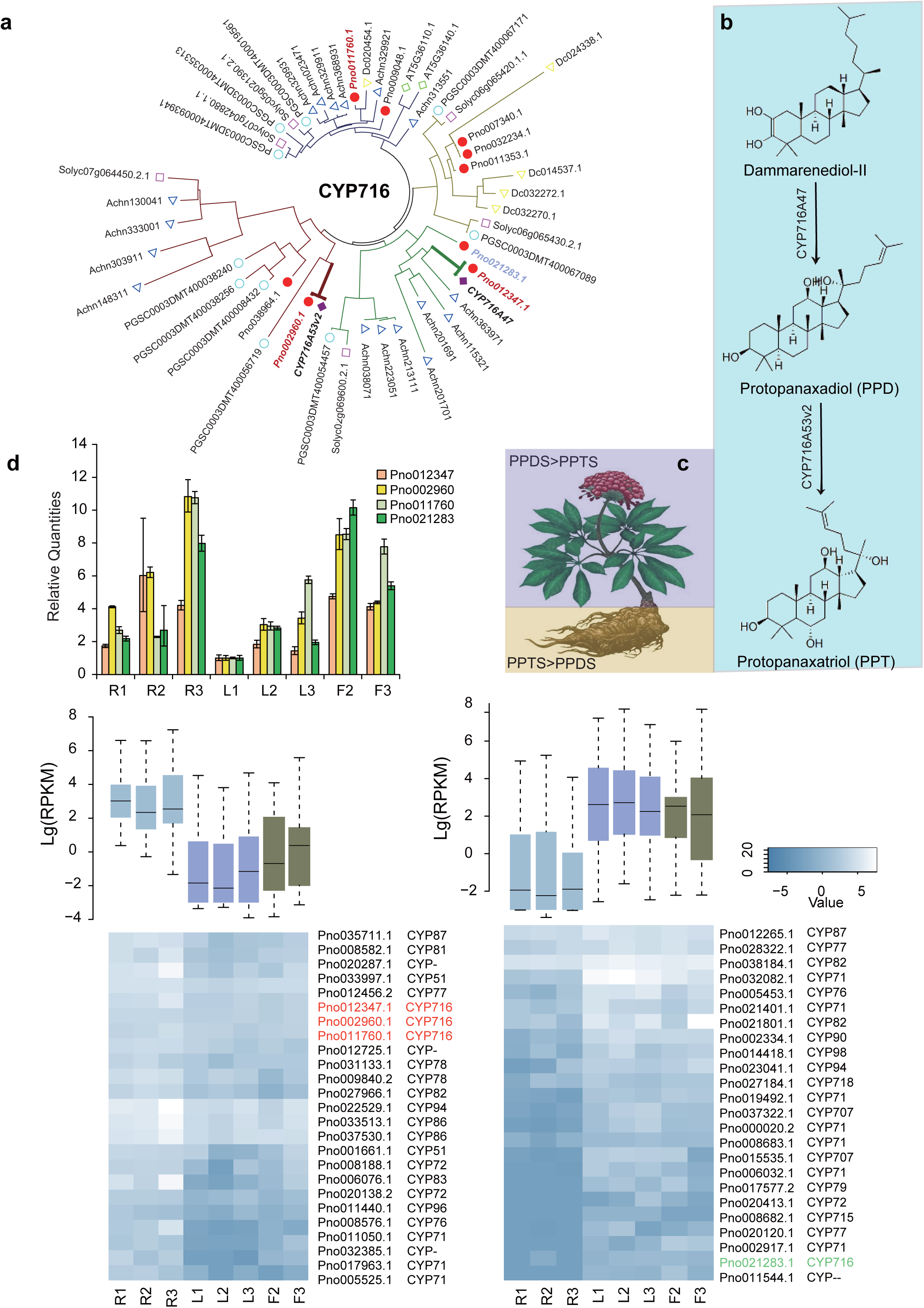
The CYP450 gene families of *P. notoginseng*. **(a)** Phylogenetic analysis of CYP716 subfamily among *P. notoginseng*, *P. ginseng* and other plants. Red circles represent *P. notoginseng*, purple diamonds represent *P. ginseng*, green circles represent *S. tuberosum*, yellow triangles represent *D. carota*, purple squares represent
*S. lycopersicum* and green diamonds represent *A. thaliana*. The lengths of clades in the gene tree show the protein similarity between two clades. **(b)** Synthesis pathway of protopanaxadiol (PPD) and protopanaxatriol (PPT) of *P. notoginseng*. **(c)**The relative amount of PPD was more than that of PPT in the aerial parts of *P. notoginseng*, but the opposite in roots. **(d)** The differentially expressed CYP450 genes of *P. notoginseng* between the aerial parts and roots and the qPCR results. The pink and green genes in the heatmap represent CYP716 subfamily genes that were involved in the synthesis pathway in *P. ginseng*.

#### UGTs

Three UGTs (*UGT71A27*, *UGT74AE2* and *UGT94Q2*) that participate in the biosynthesis of PPD-type ginsenosides and three UGTs (*UGTPg1*, *UGTPg100* and *UGTPg101*) participating in the formation of PPT-type saponins have previously been identified in *P. ginseng*. We identified a total of 160 UGT genes in the *P. notoginseng* genome based on 116 homologous UGT genes of *A. thaliana*^44^. We classified these 160 UGTs of *P. notoginseng* into 12 subfamilies containing 19 *UGT71A* homologs and 14 UTG74 homologs (**Supplementary Fig. 12**). *UGTPg1*, *UGTPg100*, *UGTPg101*, *UGTPg102* and *UGTPg103* of *P. ginseng* were *UGT71A27* homologs; however, *UGTPg102* and *UGTPg103* were found to have no detectable activity on PPT due to the lack of several key amino acids in their proteins (Fig. 4a and b). We identified 38 differentially expressed UGT genes between roots and aerial parts (flowers and leaves), including 23 expressed more highly in roots (HR) and 15 in aerial tissues (HA) (Fig. 4c and d). Interestingly, there were three *UGT71A27* homolog (Pno020280.1, Pno027722.1 and Pno026280.1) genes in the HR, and HA included four UGT74 genes (Pno031515.1, Pno033394.1, Pno002810.1 and Pno000722.1) and one *UGT71A27* homolog (Pno013844.1) gene (Fig. 4c and d). Furthermore, when comparing the *UGT71A27* homolog genes of *P. notoginseng* with *UTGPg1*, *UGTPg100*, *UGTPg101*, *UGTPg102* and *UGTPg103*, we checked the known key amino-acid residues determining the function of UGTs. We found that the key amino-acid residues *UGTPg100*-A142T, *UGTPg100*-L186S, *UGTPg100*-G338R and *UGTPg1*-H82C were conserved in both Pno026280.1 and Pno27722.1. However, *UGTPg1*-H144F is a specific mutation F in the Pno026280.1 (**Supplementary Fig. 13**). All these candidate genes probably play important roles in the biosynthesis of ginsenoside.

**Fig. 4.**
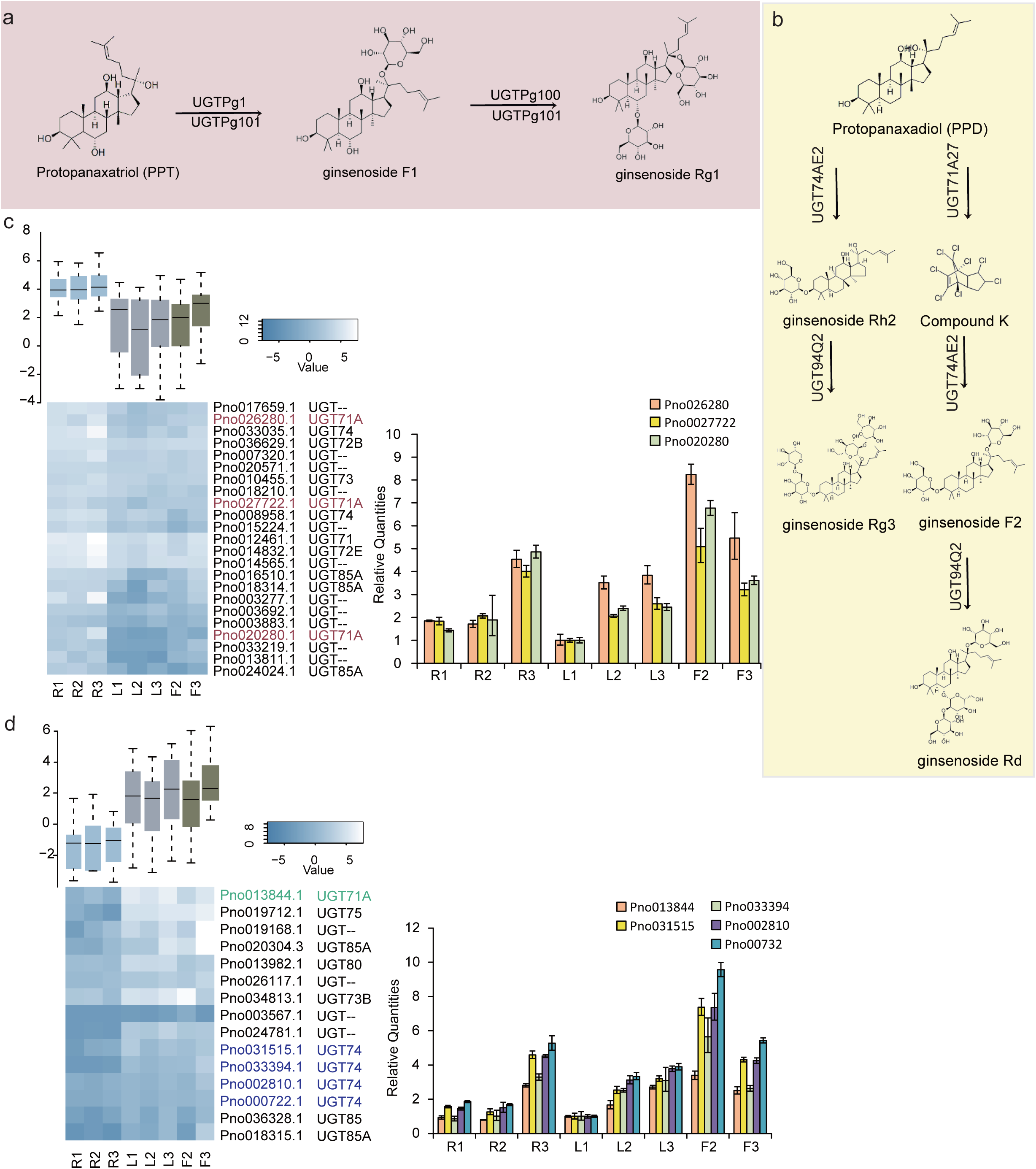
The UGT gene families of *P. notoginseng*. **(a)** The synthesis pathway of different PPT-type ginsenosides that were catalyzed by different UGT genes. **(b)** The synthesis pathway of different PPD-type ginsenosides that were catalyzed by different UGT genes. **(c)** Highly expressed UGT genes of *P. notoginseng* in roots than in aerial parts and the qPCR results. **(d)** Highly expressed UGT genes of *P. notoginseng* in aerial parts than in roots and the qRCR results. The purple, green and blue genes in the heatmap represent gene families that were involved in the synthesis pathway in *P. ginseng*.

### Identification of transcription factor binding sites

We have predicted the transcription factor binding site (TFBS) in the upstream regions of ten genes encoding enzymes involved in the ginsenoside biosynthesis, including HMG-CoA synthase (HMGS), HMG-CoA reductase (HMGR), mevalonate kinase (MK), phosphomevalonate kinase (PMK), mevalonate diphosphate decarboxylase (MDD), isopentenylpyrophosphate isomerase (IPI), geranylgeranyl diphosphate synthase (GGR), farnesyl diphosphate synthase (FPS), squalene epoxidase (SE) and dammarenediol-II synthase (DS) (**Supplementary Fig. 14**). DNA-binding with one finger 1 (Dof1) and prolamin box binding factor (PBF) binding sites were detected in all these ten genes which used the same binding site motif (5′-AA[AG]G-3′) (**Supplementary Table 13**). Therefore, the expression levels of Dof1 and PBF with these ten genes in different developmental stages of roots and leaves from *P. notoginseng* were compared (**Supplementary Table 14**). Most of genes’ expression levels were higher in the 3 roots than that in the 3 leaves, expecting PMK and Dof1 (**Supplementary Fig. 15**). Dof1 belong to the Dof family, which is plant-specific transcription factor ^45^. Yanagisawa’s studies suggested that Dof1 in maize is related with the light-regulated plant-specific gene expression ^46,47^. PBF encodes Dof zinc finger DNA binding proteins, which could enhance the DNA binding of the bZIP transcription factor Opaque-2 to O2 binding site elements ^48^. In three stages of roots, the trend of PBF expression level was consistent with that of DS, HMGR, SE, HMGS, MK, MDD, FPS and GGR (**Supplementary Table 15**). Whereas, in different developmental stage of leaves, there was almost no significant change in genes expression levels in these genes, excluding Dof1 and DS with a rising trend (**Supplementary Table 16**). We inferred that the PBF and Dof1 were probably associated with the transcriptional controls of these genes, involved in triterpene saponins biosynthesis pathway, in different parts, the root and the leaf of *P. notoginseng*, respectively.

## Discussion

Compared with other phylogenetically-related plant species, there was notable gene family contraction of R-genes in *P. notoginseng.* The encoded resistance proteins play important roles in plant defense by controlling the host plant’s ability to detect a pathogen attack and facilitate a counter attack against the pathogen^5^. Whether *P. notoginseng* has evolved other defenses and/or anti-stress mechanisms to compensate for the contraction of R-genes remains to be determined in data mining of the first draft genome of this genus.

Given that the notable contraction of R-genes of *P. notoginseng* may provide clues to its poor stress and disease resistance, we also observed expansion and contraction of different subfamilies of the ABC transporter gene family in *P. notoginseng*. The substrates of ABC proteins include a wide range of compounds such as peptides, lipids, heavy metal chelates and steroids^49^. Interestingly, one of the contracted gene candidates (Pno003332.1) in *P. notoginseng* was reported to be involved in iron export^35^. A previous experimental study showed that optimal iron concentration could significantly determine the plant biomass and stabilize arsenic content in soil to reduce arsenic contamination^50^ where arsenic contamination of *P. notoginseng* is a serious agricultural problem due to arsenic pollution in the environment^50^.

Phenolics, alkaloids and terpenoids are three major classes of chemicals involved in plant defenses^51^. Another possible complementary mechanism to cope with reduced pathogen resistance, which possibly resulted from R-gene contraction, is through evolving synthesis of defensive secondary plant metabolites. Pepper and tomato, two species phylogenetically close to *P. notoginseng*, produce alkaloids such as capsaicinoids and tomatine, respectively, which have been found to function as deterrents against pathogens^20,52^. Azadiracht, a meliacane-type triterpenoid with very similar skeleton to dammarane from the neem tree (*Azadirachta indica*)^53^, has insecticidal properties. Ginsenosides are the oligosaccharide glycosides of a series of dammarane- or oleanane-type triterpenoid glycosides^9^. Palazón *et al.* reported that during *in vitro* culture of ginseng hair root lines, the addition of methyl jasmonate, a signaling molecule specifically expressed by plants in response to insect and pathogenic attacks, could enhance overall ginsenoside production and conversion of PPD-type (e.g. Rb1, Rb2, Rc and Rd) to PPT-type (e.g. Re, Rf and Rg1) ginsenoside^54^.

In addition, natural ginsenosides have antimicrobial and antifungal action, as shown in numerous laboratory studies^55,56^. The exact natural roles of ginsenosides in plants, particularly controlling plant growth and defense against pathogenic infection, remain unclear and require further investigation.

*DDS* is the critical determinant for the initial step of producing the dammarane-type sapogenin (aglycone of saponin) template. This can be further subjected to structural diversification by *CYP450* and UGT enzymes. Along the triterpenoid saponin biosynthesis pathway, DDS of the OSC group is responsible for the cyclization of 2,3-oxidosqualene to dammarenediol-II^57^. A highly-conserved *DDS* homolog of *P. ginseng* was also identified in *P. notoginseng*. Upstream of *DDS* in the triterpenoid saponin biosynthesis pathway, three *SQLE* and two *FDFT1* genes were identified and shown to exhibit significant differential expression in different parts in *P. notoginseng*.

Han *et al.* identified that *CYP716A47* produces PPD through the hydroxylation of C-12 in dammarenediol-II^43^, *CYP716A53v2* hydroxylates the C-6 of PPD to produce PPT^58^, and *CYP716A52v2* is involved in oleanane-type ginsenoside biosynthesis^59^ in *P. ginseng*. We were interested in determining how the transcriptional regulation of different CYP450 genes respectively correlated with synthesis of higher content of PPT- and PPD-type ginsenosides in roots and aerial parts (leaves and flowers). Combined the previous research and transcriptome analysis, we identified the key genes which produces PPD and PPT in the *P. notoginseng*.

UGTs are GTs that use uridine diphosphate (*UDP*)-activated sugar molecules as donors. Several types of UGTs, which catalyze the synthesis of specific sapogenins to ginsenosides, were found in previous studies of *P. ginseng*^60^. For example, UGTPg1 was demonstrated to region-specifically glycosylate C20-OH of PPD and PPT^61^. *UGTPg100* specifically glycosylates *PPT* to produce bioactive ginsenoside Rh1, and *UGTPg101* catalyzes *PPT* to produce F1^61^. Up to now, very few of the UGTs characterized that are able to glycosylate triterpenoid saponins have been found in *P. ginseng*^62-65^ and *P. notoginseng*^66,67^. However, in the present study, an almost complete collection of 160 UGT genes, representing many new homologs and putative UGT members, was disclosed in the *P. notoginseng* genome. In our study, we found the lack of correlation between qPCR and RNA-seq in the Pno026280, Pno002772 and Pno020280. In order to eliminate error because of improper normalization method, we used other two quantification methods. Using the raw RNA-seq reads counting and normalized by quantiles (limma) ^68^and DESeq^69^ showed the completely consistent tendency with FPKM results (**Supplementary Table 17**). The probable reasons are the nonspecific binding of qPCR primers to cDNA template and the bias in the RNA-seq sequencing due to sequencing library preparation, namely, it was due to nature of these two methods (RNA-seq vs. qPCR).

In conclusion, using a combination of data from genome and transcriptomes, we identified contraction and expansion in numerous important genome events during the evolution of this plant. The contraction of disease-resistance genes families figured out the vulnerability to disease of *P. notoginseng* on genetic level and the expansion of several *ABC* transporter subfamilies indicated the potential of *ABC*s as an additional mechanism for the plant to cope with biotic stress. Combining eight transcriptomes of roots and aerial parts, several key genes and all their family members involved in the synthesis pathway of ginsenosides were also identified including dammarenediol synthase, *CYP716* and *UGT71*. These findings provide new insight into its poor resistance and the biosynthesis of high-valued pharmacological molecules. Finally, it is essential that a better chromosome level reference genome assembly using long reads sequencing platform and Hi-C technology, serving as an important platform for improving breeding and cultivation of this vulnerable plant species to increase the medicinal and economic value of *Panax* species.

## Methods

### K-mer analysis

We used the total length of sequence reads divided by sequencing depth to calculate the genome size. To estimate the sequencing depth, the copy number of 17-mers present in sequence reads was counted and the distribution of sequencing depth of the assembled genome was plotted. The peak value in the curve represents the overall sequencing depth, and can be used to estimate genome size (G): G = Number_17-mer_/Depth_17-mer_.

### Genome assembly

The first genome assembly version (v1.0) using *SOAPdenovo^70^* (v2.04) based on the short length reads data sequencing by Hiseq2000, was highly fragmented. It contained more than 600Mb gaps with missing sequence because of presence of high proportion of repetitive DNA sequences in the genome. The contig N50 sizes of v1.0 was 4.2kb. For the short contig N50 size, therefore, we further generated ~13Gb SMRT sequence data (~6-fold of whole genome size) with average sub-read length of ~9kb. Considering the inadequate depth of our Pacbio third generation sequencing, we used the PBJelly^71^ to fill the gap and upgrade the genome assembly with long length reads. Concretely, we generated our protocol file containing full paths of the reference assembly above, the output directory and the input files. Then, we executed multiple steps (including setup, mapping, support, extraction, assembly, output) in a consecutive order. From the statistics of the result of PBJelly, we found that the gap number was lowered to 533Mb while sizes of contig N50 was increased to 10.7kb. Next, we used the libraries with the insert sizes of 500bp and 800bp as the gap-filling input data of the GapCloser (v1.0, http://soap.genomics.org.cn/). After running, the gap number was decreased to 493Mb and the length of contig N50 was increased to 16.4kb. Finally, the sizes of contig and scaffold N50 of the final version (v1.1) were 16.4kb and 70.6kb, respectively. In addition, the total length of the genome changed from 2.1Gb to 2.4Gb with 493Mb gap sequences. The detailed parameters of *SOAPdenovo* were “pregraph -s lib.cfg.1 -z 2800000000 -K 45 -R -d 1 -o SAN_63 -p 12; contig -g SAN_63 -R -p 24; map -s lib.cfg.1 -g SAN_63 -k 45 -p 24; scaff -g SAN_63 -F -p 24”. We also adopted ABySS^72^ to estimate the optimized K value for genome assembly from 35 to 75 using about 20-fold high quality sequencing reads. The result revealed the K=45 is the best K value (**Supplementary Table 18**), which is completely consistent with the best K value of SOAP*denovo*.

### Transcript *de novo* assembly

Leaves and roots of *P. notoginseng* were collected from 1-, 2- and 3-year-old *P. notoginseng* and flowers collected from 2- and 3-year-old *P. notoginseng*. All of these 8 samples were randomly collected from the fileld of commercial planting base in Yanshan county of Wenshan Zhuang and Miao minority autonomous prefecture in Yunnan province. After cleansing, the leaves, roots and flowers were collected separately, cut into small pieces, immediately frozen in liquid nitrogen, and stored at −80 °C until further processing. Subsequently, mRNA isolation, cDNA library construction and sequencing were performed orderly. Briefly, total RNA was extracted from each tissue using TRIzol reagent and digested with DNase I according to the manufacturer’s protocol. Next, Oligo magnetic beads were used to isolate mrna from the total rna. By mixing with fragmentation buffer, the mrna was broken into short fragments. The cDNA was synthesized using the mRNA fragments as templates. The short fragments were purified and resolved with EB buffer for end repair and single nucleotide A (adenine) addition, and then connected with adapters. Suitable fragments were selected for PCR amplification as templates. During the quality control steps, an Agilent 2100 Bioanalyzer and ABI StepOnePlus Real-Time PCR System were used for quantification and qualification of the sample library. Each cDNA library was sequenced in a single lane of the Illumina HiSeqTM 2000 system using paired end protocols. Before performing the assembly, firstly, raw sequencing data that generated by Illumina HiSeq2000, was subjected to quality control (QC) check including the analysis of base composition and quality. After QC, raw reads which contained the sequence of adapter (adapters are added to cDNAs during library construction and part of them may be sequenced), more than 10% unknown bases or 5% low quality bases were filtered into clean reads. Finally, Trinity^73^ was used to perform the assembly of clean reads (detail information could be retrieved in Trinity website http://trinityrnaseq.sourceforge.net/). Based on the Trinity original assembly result, TGICL^74^ and Phrap^75^ were used to acquire sequences that cannot be extended on either end, the sequences are final assembled transcripts. We mapped all transcripts that were *de novo* assembled from nine transcriptome short reads by Trinity into the genome assembly using Blat^76^, and then calculated their coverage and mapping rate. The sequencing depth distribution was examined by aligning all short reads against assembly using SOAP2^77^ and then the depth of each base was calculated.

### Gene expression analysis

All the QC and filter processes of raw sequencing transcriptome data were the same as that of *de novo* assembly. All the clean reads are mapped to reference gene sequences by using *SOAP*2^77^ with no more than 5 mismatches allowed in the alignment. We also performed the QC of alignment results because mRNAs were firstly broken into short segments by chemical methods during the library construction and then sequenced. If the randomness was poor, read preference for specific gene region could influence the calculation of gene expression. The gene expression level of each gene was calculated by using RPKM method^78^ (reads per kilobase transcriptome per million mapped reads) based on the unique alignment results. In the previous test, comparative analysis between RPKM and FPKM showed that the correlation between RPKM and QPCR was better than the correlation between FPKM and QPCR. Referring to the previous study^79^, we have developed a stringent algorithm to identify differentially expressed genes between two samples. The probability of a gene expressed equally between two samples was calculated based on the Poisson distribution. P-values corresponding to differential gene expression test were generated using Benjamini, Yekutieli.2001 FDR method^80^. Moreover, because of some debates in the normalization method by gene size like RPKM and then two new quantification methods were conducted to validate the existing results. Concretely, based on the tophat mapping results, R package ‘Rsubread’ was used to do the raw reads counting then we used ‘limma’ with the normalize parameter “quantile”. Additionally, DESeq also was conducted. Finally, aiming at the differential expressed genes mentioned in the article based on RNA-seq (RPKM), we found the completely same tendency.

### Identification of repetitive sequences

Transposable elements (TEs) in the *P. notoginseng* genome were annotated with structure-based analyses and homology-based comparisons. In detail, RepeatModeler^22^ was used to *de novo* find these TEs based on features of structures.

RepeatMasker and RepeatProteinMask were applied using RepBase^81^ for TE identification at the DNA and protein level, respectively. Overlapped TEs belonging to the same repeat class were checked and redundant sequences were removed. TRF software^82^ was applied to annotate tandem repeats in the genome.

### Gene prediction and annotation

We predicted protein-encoding genes by homolog, *de novo* and RNA-seq evidence; and results of the former two methods were integrated by the GLEAN program^83^ then filtered by threshold level of 20% percent overlap with at least 3 homolog species support. Subsequently, the inner auto pipeline was used to integrated RNA-seq data within and then checked manually. Firstly, protein sequences from five closely related species—*A. thaliana*, *S. lycopersicum*, *S. tuberosum*, *Capsicum annuum* and *Daucus carota*—were applied for homolog prediction by respectively mapping them to the genome assembly using TblastN software with E-value −1e-5. Aiming at these target areas, we used Genewise^84^ to cluster and filter pseudogenes. Then Fgenesh^85^, AUGUSTUS^86^ and GlimmerHMM^87^ were used for *de novo* prediction with parameters trained on A. thaliana. We merged three de novo predictions into a unigene set. *De novo* gene models that were supported by one more *de novo* methods were retained. For overlapping gene models, the longest one was selected and finally, we got de novo-based gene models. Using *de novo* gene set (30,660-41,495) and five homolog-based results as gene models (24,358 to 36,116) integration was done using the GLEAN program. Finally, we got the GLEAN gene set (referred to as G-set, 31,678) with parameter “value 0.01, fixed 0”. Combined with the RNA transcriptome sequencing data, we used Tophat2^88^ to map and Cufflinks^89^ to assemble reads into transcripts and finally integrated the transcripts into the gene set using inner pipeline. The process of inner pipeline was: (1) The assembly of transcriptome data. RNA-seq reads were mapped to *P. notoginseng* genome using tophat, and then cufflinks pipeline was used to conduct the assembly of transcripts. (2) Prediction of the ORF in transcripts (CDS length >=300bp, score >-15). (3) Comparison of transcripts and Glean results to count the overlap region (identity >=95%, total/len >=90%). (4) Integration of the existing results. Aiming at the results, we checked manually by discarding genes without any functional annotation information, any homolog support or cufflinks results support. The final gene set (35,451) was obtained. Then, with the known protein databases (SwissProt^90^, Trembl, KEGG^91^, InterPro^92^ and GO^93^), we annotated these functional proteins in the gene set by aligning with template sequences in the databases using BLASTP (1e-5) and InterProScan.

### Identification of R-genes

Most R-genes in plants encode NBS-LRR proteins. According to the conservative structural characteristics of domains, we used HMMER (V3, http://hmmer.janelia.org/software) to screen the domains in the Pfam NBS (NB-ARC) family. Then we compared all NBS-encoding genes with TIR HMM (PF01582) and LRR 1 HMM (PF00560) data sets using HMMER (V3). For the CC domains, we used the MARCOIL program^94^ with a threshold probability of 0.9 and double-checked using paircoil2^95^ with a P-score cut-off of 0.025.

### Gene cluster analysis

We used OrthoMCL^96^ to identify the gene family and so obtained the single-copy gene families and multi-gene families that were conserved among species. Then we constructed a phylogenetic tree based on the single-copy orthologous gene families using PhyML^97^. The different molecular clock (divergence rate) might be explained by the generation-time hypothesis. Here, we used the MCMCTREE program^98^ (a BRMC approach from the PAML package) to estimate the species divergence time. ‘Correlated molecular clock’ and ‘JC69’ model in the MCMCTREE program were used in our calculation.

### Analysis of key gene families

The reference genes of interest in *A. thaliana* (such as ABC, CYP450 and UGT) were found in the TAIR10 functional descriptions file. Then, the target genes of *P. notoginseng* were identified and classified using BlastP with cut-off value of 1e-5 and constructing gene trees using PhyML (-d aa -b 100). The tree representation was constructed using MEGA software.

### Real-time PCR analysis

Isolated RNA from different *P. notoginseng* tissues were reverse-transcribed to single-strand cDNA using the Super Script™ III First-Strand Synthesis System (Invitrogen™, USA). Quantitative reactions were performed on the Real-Time PCR Detection System (VIIA 7^TM^ Real-Time PCR System, Applied Biosystems, USA) using FastStart Universal Probe Master and Universal ProbeLibrary Probes (Roche, Switzerland). All primers used in this study are listed in **Supplementary Table 19**. The relative gene expression was calculated with the 2^-ΔΔCT^ method. For each sample, the mRNA levels of the target genes were normalized to that of 18S rRNA.

## Availability of supporting data and materials

The genome assembly and sequencing data have been deposited into NCBI Sequence Read Archive (SRA) under project number PRJNA299863 and into *Giga*DB.

## Declarations

### Acknowledgements

This work was supported by the Science and Technology Development Fund (FDCT) of Macao SAR (Ref. No. 134/2014/A3), Research Committee of University of Macau (MYRG139(Y1-L4)-ICMS12-LMY and MYRG2015-00214-ICMS-QRCM), the Overseas and Hong Kong, Macau Young Scholars Collaborative Research Fund by the Natural National Science Foundation of China (Grant no. 81328025), Technology Innovation Program Support by Shenzhen Municipal Government (NO.JSGG20130918102805062) and Basic Research Program Support by Shenzhen Municipal Government (NO.JCYJ2014072916361729 and NO. JCYJ20150831201123287).

## Additional files

**Additional file 1: Supplementary information** includes supporting figures and supporting tables.

## Author’s contributions

S. M.Y.L, X.X and X.L designed the project. G.F, B.R.Y, Y. F, H.Z, X.L, C.S, K.M, J.H, R.G, L.S, S.D, Q.X, W.L, M.L, A.K.W, D.Z and M.C analyzed the data. W.C, X.L, G.F, Y. F, J.C and B.R.Y wrote the manuscript. B.R.Y, M.G and C. H prepared the samples and conducted the experiments.

## Authors’ Affiliations

(1) State Key Laboratory of Quality Research in Chinese Medicine, Institute of Chinese Medical Sciences, University of Macau, Macao, China.

(2) BGI-Shenzhen, Shenzhen 518083, China.

(3) State Key Laboratory of Bioelectronics, School of Biological Sciences and Medical Engineering, Southeast University, Nanjing 210096, China

(4) Faculty of Science and Technology, Department of Civil and Environmental Engineering, University of Macau, Macao, China

